# MicroRNA-195-3p as a potential regulator of hypoxic injury in HUVECs

**DOI:** 10.1101/2023.02.23.529661

**Authors:** Wenjing Zhang, Bingshi Liu, Yanfang Wang, Lixian Sun, Chao Liu, Haoran Zhang, Wei Qin, Jingyi Liu, Leng Han, Zhoukai Cui, Weichao Shan

**Affiliations:** Department of Cardiology, Affiliated Hospital of Chengde Medical University, Chengde, China; Department of Cardiology, Pingquan City Hospital, Chengde, China

**Keywords:** miR-195-3p, atherosclerosis, human umbilical vein endothelial cells (HUVECs), hypoxia injury, proliferation, migration, apoptosis, autophagy

## Abstract

Coronary heart disease is the most common heart disease and is the leading cause of cardiovascular death worldwide. Revascularization methods are considered effective against coronary heart disease However, the mechanism of molecular revascularization remains largely unknown. Endothelial cells are the primary cells that initiate angiogenesis and arteriogenesis and require a hypoxic environment for induction. In this study, we aimed to determine the expression and role of microRNA-195-3p in hypoxia-treated HUVEs(human umbilical vein endothelial cells). Herein, we induced hypoxia in human umbilical vein endothelial cells using the “Anaerobic tank method.” Hypoxia injured human umbilical vein endothelial cells showed upregulation of microRNA-195-3p; decreased cell proliferation, migration, and autophagy; and increased apoptosis. Furthermore, the microRNA-195-3p inhibitor partially reversed the effects of hypoxia-induced injury of human umbilical vein endothelial cells. Therapeutic intervention using microRNA-195-3p inhibitor could maintain endothelial cell function under hypoxic conditions, improve cell activity, and be considered a new treatment strategy for coronary heart diseases.

## INTRODUCTION

Coronary heart disease (CHD) and its cardiovascular sequelae represent the leading cause of mortality worldwide. Despite significant advances in revascularization and intensive medical care interventions, patients with CHD who undergo percutaneous transluminal angioplasty have a persistently high rate of myocardial infarction and death(1).Clinical studies have identified triggered angiogenesis as an effective therapy for CHD(2). Endothelial cells (ECs) are the primary cells that undergo activation and initiate arteriogenesis, resulting in the formation of collateral circulation(3,4). Therefore, it is essential to identify specific biomarkers that regulate the proliferation and apoptosis of microvascular ECs and determine their underlying mechanism.

MicroRNAs (miRNAs) are endogenous non-coding RNAs approximately 22 nucleotides long that regulate the expression of specific target mRNAs; increasing studies have confirmed that miRNAs play essential roles in regulating cardiac functions(5–8). miR-195 belongs to the miR-15/107 family and are stress-inducible miRNAs activated in multiple diseases, including cardiovascular diseases(9). Increasing miRNA-195 levels in failing myocardium are associated with metabolic dysfunction and cardiomyopathy(10). In contrast, miR-195 promotes apoptosis in cardiomyocytes in the diabetes model via downregulating Bcl-2 and Sirt1/2(11). miR-195-3p and miR-195-5p are two mature forms of miR-195(12). Several studies have revealed the role of miR-195-3p in cardiovascular diseases, but its role in hypoxia-induced ECs remains unidentified.

Human umbilical vein endothelial cells (HUVECs) are widely used to study vascular diseases and endothelial dysfunction(13). In this study, we aimed to investigate the expression and role of miR-195-3p in hypoxia-treated HUVECs. We found an upregulation of miR-195-3p in hypoxia-cultured HUVECs. Inhibition of miR-195-3p restored hypoxia-induced reduced cell proliferation, migration, and autophagy and inhibited hypoxia-induced apoptosis of HUVECs. Therefore, targeting miR-195-3p may have potential benefits for CHD therapy.

## MATERIALS AND METHODS

### Cell culture and hypoxia induction

The present study was approved by the Ethics Committee of the Affiliated Hospital of Chengde Medical University. HUVECs (Beijing Being Chuanglian Biotechnology Institute, Beijing, China) were cultured in Dulbecco’s Modified Eagle’s Medium (DMEM,Gibco,USA) supplemented with 1% penicillin and streptomycin, 10% fetal bovine serum (FBS,Gibco,China), and incubated at 37°C with 5% CO_2_. Cells in the logarithmic phase were collected for use in subsequent experiments.

HUVECs were treated with “Anaerobic tank method” (14) to develop hypoxia-induced ECs injury in vitro. Briefly, HUVECS were cultured in low-glucose DMEM without FBS at 37°C in an incubator maintained at 95% N_2_ and 5% CO_2_ for 6 h. Following the successful establishment of the hypoxia model, the cells were used for subsequent experiments.

### Western blotting

The protein expression levels were analyzed using western blotting. Total protein was quantified, and 30 μg protein per lane was separated using 8% Sodium dodecyl-sulfate polyacrylamide gel electrophoresis and transferred to a membrane. The membranes were then incubated with the following primary antibodies at 4°C for 6 h: GAPDH (1:6000, no. A19056), vinculin (1:1000, A2752) (ABclonal Technology, Co., China), caspase-3 (1:1000, no. #14220), cleaved caspase-3 (1:1000, no. #9664), LC3B (1:1000, no. #3868), Cytochrome C (1:1000, no. #11940), hypoxia inducible factor 1 subunit alpha (HIF1A; 1:1000, no. #14179), Bax (1:1000, no. #2772), and Bcl-2 (1:1000, no. #4223) (Cell Signaling Technology, Inc., USA). Following primary antibody incubation, the membranes were incubated with secondary antibody(Goat Anti-rabbit IgG, Abcam, ab182016) at room temperature for 2 h. Protein bands were visualized using enhanced chemiluminescence reagent. The bands were quantified using ImageJ. GAPDH or vinculin was used as the internal reference protein.

### miRNA inhibitor transfection

The miR-195-3p inhibitor (5’-GGAGCAGCACAGCCAAUAUUGG-3’) was a single-stranded antisense RNA against miRNAs with 2’-O-Me modification. HUVECs were transfected with a mixture containing 10 μL (50 nM) inhibitor, 10 μL Lipofectamine 3000, and 980 μL Opti-MEM in DMEM with 1% FBS. Cells transfected with negative control (NC) inhibitors were used as controls. HUVECs were harvested and lysed for polymerase chain reaction or western blot 48 h after transfection.

### Quantitative reverse transcription-polymerase chain reaction

A miRcute (Tiangen, China) was used to extract total RNA. Primers were designed and synthesized by TIANGEN BIOTECH CO., LTD (Tiangen, China). U6 primer sequence: Forward, 5’-CTGGCTTCGGCAGCACA-3’; Reverse, 5’-AACGCTTCACGAATTTGCGT-3’. Primer sequence for has-miR-195-3p: Forward, 5’-CCAAUAUUGGCUGUGCUGCUCC-3’. RNA was reverse transcribed into cDNA using cDNA reverse transcription kit (Tiangen, China) following manufacturer’s instructions. With U6 serving as the internal reference, the relative transcription level of the target genes was calculated using the 2^-ΔΔCt^ method. The ΔΔCt was calculated following the equation:

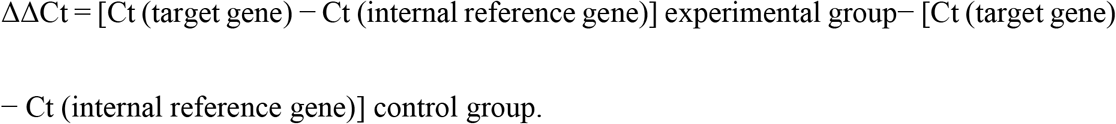

### 5-ethynyl-2’-deoxyuridine assay

Cells were seeded in 48-well plates for 24 h and incubated in medium containing 5-ethynyl-2’-deoxyuridine (EdU) (50 mM; Guangzhou RiboBio Co., Ltd.) for 2 h. Cell fixation, staining, and DNA staining were performed according to the instructions. The nucleus and EdU-positive cells were observed under a fluorescence microscope (Nikon, Tokyo, Japan), and the proliferation rate (EdU-positive cells/nucleus) at random fields were calculated.

### Colony formation assay

Cells were plated in 6-well plates (2000 cells/well) and incubated in DMEM with 10% FBS at 37°C. The cells were fixed and stained with 0.1% crystal violet two weeks later. The number of visible colonies was counted manually.

### Cell counting kit-8 assay

HUVECS viability was assessed by Cell Counting Kit-8 (CCK-8, Dojindo, Japan) assay as described previously. In brief, the cell densities were 1 × 10^3^ cells/well for CCK-8. After being treated, 90 mL of DMEM containing 10 mL of CCK-8 solution was added to each well, and the 96-well plates were incubated for 2 h at 37°C. The absorbance was determined with a spectrophotom eter at 450 nm (BioTek, Winooski, VT, USA)

### Wound healing assay

Cells were seeded into 6-well plates, grown to confluence, and incubated in a serum-free medium (SFM) for 4 h. Afterward, a sterile 200 mL plastic pipette tip was used to create a wound in the center of the well. SFM was then replaced by DMEM containing 1% FBS. Images were taken at 0 and 24 h using a microscope, and the results were analyzed using ImageJ software.

### Transwell assay

A total of 1×10^6^ cells were seeded into the upper chamber of the Transwell in DMEM, while the lower chamber was maintained in a 10% FBS medium. After 8 h of incubation, cells at the bottom layer of the upper chamber were fixed using 4% paraformaldehyde for 30 min, stained with 0.5% crystal violet (Solarbio, China) for 15 min, and counted under an inverted microscope.

### Terminal deoxynucleotidyl transferase dUTP nick end labeling assay

HUVECs were cultured on a 24-well plate, transfected with 50 nmol/L miR-195-3p inhibitor/inhibitor NC control, and stimulated with hypoxia for 6 h. The cells were stained using terminal deoxynucleotidyl transferase dUTP nick end labeling (TUNEL apoptosis assay kit, Roche, American), and the cells were viewed under a fluorescence microscope.

### Statistical analysis

The data were analyzed using GraphPad Prism software 8.0. Data were calculated as mean ± standard deviation. A two-sided student t-test was used to analyze individual differences; p < 0.05 was considered statistically significant.

## RESULTS

### Effects of hypoxia induction on miR-195

The expression of miR-195 in hypoxia-induced HUVECs was detected. Compared with the control group, no significant difference in the expression of miR-195-5p was observed after 6 h of hypoxia induction in HUVECs. In contrast, the expression of miR-195-3p was significantly increased (p < 0.0001) (Figure 1 · A, B), suggesting the role of miR-195-3p in the hypoxic injury of HUVECs.

**FIGURE 1.**
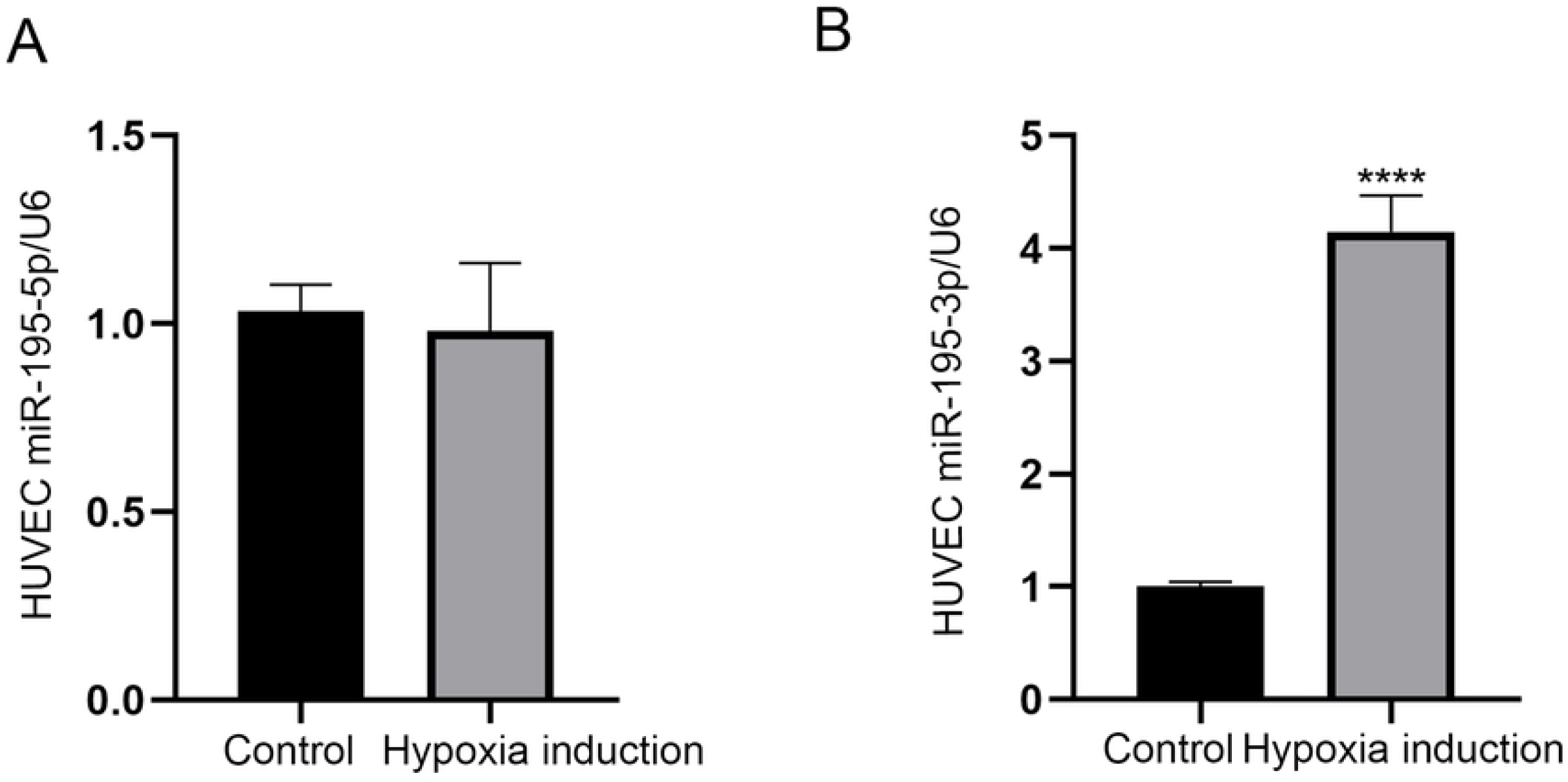
miR-195-3p is involved in the hypoxia-mediated injury of HUVECs. (**A**) No difference in miR-195-5p following 6 h of hypoxia induction compared to the control group. (**B**) The expressions of miR-195-3p increased following 6 h of hypoxia induction in HUVECs compared to the control group. All the values are the mean of three independent experiments (n=3). ***p < 0.001 vs. control. HUVECs, human umbilical vein endothelial cells.

### miR-195-3p inhibitor in the hypoxic injury of HUVECs

To clarify whether miR-195-3p is involved in the hypoxia injury, a miR-195-3p inhibitor was used. The miR-195-3p inhibitor was transfected to HUVECs using liposome-mediated transfection. Treatment with the inhibitor reduced the gene expression of miR-195-3p compared with the NC(p < 0.01) (Figure 2). These results imply the successful transfection of the miR-195-3p inhibitor into the HUVECs.

**FIGURE 2.**
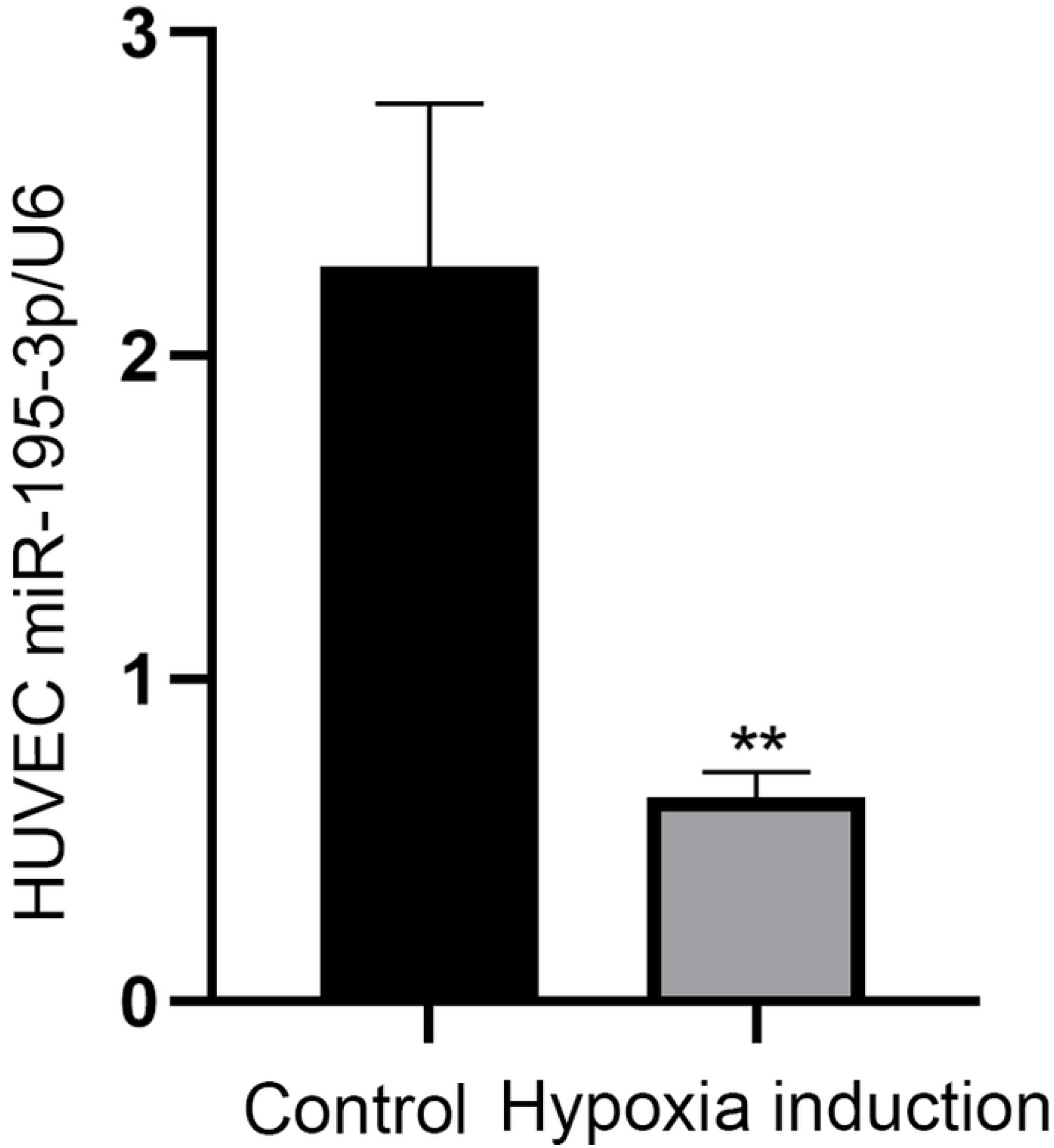
Effects of miR-195-3p inhibitor on HUVECs. The expression of miR-195-3p in miR-195-3p inhibitor treatment group. All the values are the mean of three independent experiments (n=3). **p < 0.01 vs. control. HUVECs, human umbilical vein endothelial cells.

### miR-195-3p inhibitor partially restores the proliferation in damaged HUVECs

The inhibitor of miR-195-3p was transfected to the HUVECs, and hypoxia was induced for 6 h. The cells were then analyzed for their proliferation rate and cell viability. Treatment with an inhibitor following hypoxia treatment restored the EdU-positive cells comparable to the control group (p < 0.001) (Figure 3A, B). Similarly, the CCK-8 assay showed that transfection with miR-195-3p inhibitor improved the cell viability of hypoxia-injured HUVECs, confirming a partial restoration of cell proliferation(p < 0.001) (Figure 3C). The miR-195-3p inhibitor also enhanced colony-forming ability (p < 0.001) (Figure 3D, E). In summary, the miR-195-3p inhibitor reversed hypoxia-induced proliferation inhibition.

**FIGURE 3.**
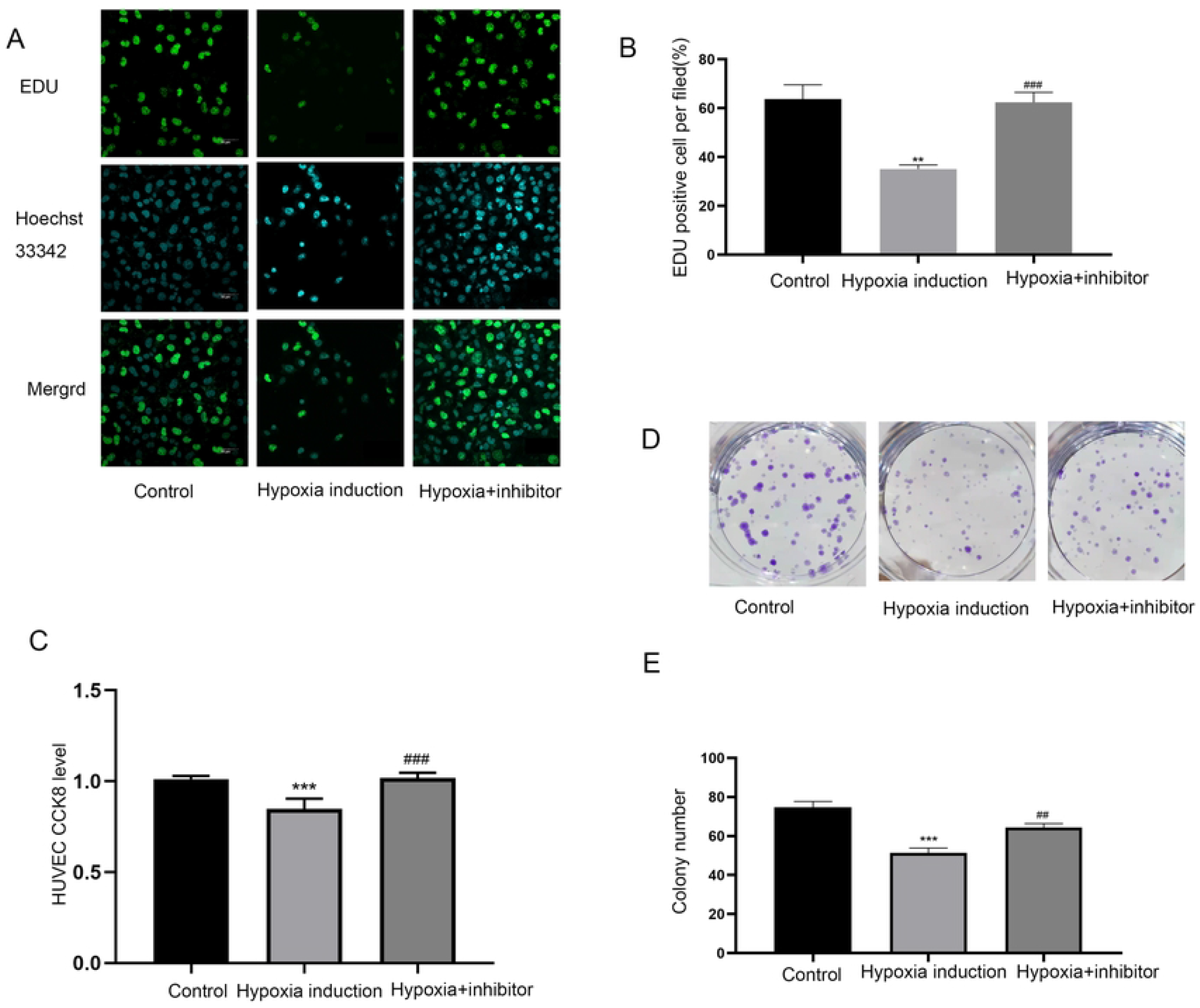
miR-195-3P inhibitor regains hypoxia-induced HUVECs proliferation rate. (**A, B**) EdU results shows the proliferation rate of each group. The cells were stained with Hoechst 33342 (5 μg/mL), and the proliferation population was analyzed (n=3). Scale bar, 50 μm. (**C**) The viabilities of cells in the different treatment groups was measured using the CCK-8 assay (n=5). (**D, E**) Representative images of cell colonies. Colony formation assays determined cell proliferation in HUVECs. **p < 0.01, ***p < 0.001 vs. the control group. ##p < 0.01, ###p < 0.001 vs. hypoxia induction for 6 h group. HUVECs, human umbilical vein endothelial cells; EdU, 5-ethynyl-2’-deoxyuridine; CCK-8, Cell Counting kit-8.

### miR-195-3p inhibitor enhanced the migration ability of hypoxia injured HUVECs

The scratch-wound healing and Transwell experiments were used to study the cell migration level of HUVECs. In the control group, hypoxia induction for 6 h reduced the migration of HUVECs. However, downregulation of miR-195-3p reversed this result. HUVECs transfected with miR-195-3p inhibitor and exposed to hypoxia for 6 h showed enhanced wound healing (p < 0.01) (Figure 4A, B) and Transwell (p < 0.01) (Figure 4C, D) migration properties. The miR-195-3p inhibitor can improve the migration ability of HUVECs damaged by hypoxia.

**FIGURE 4.**
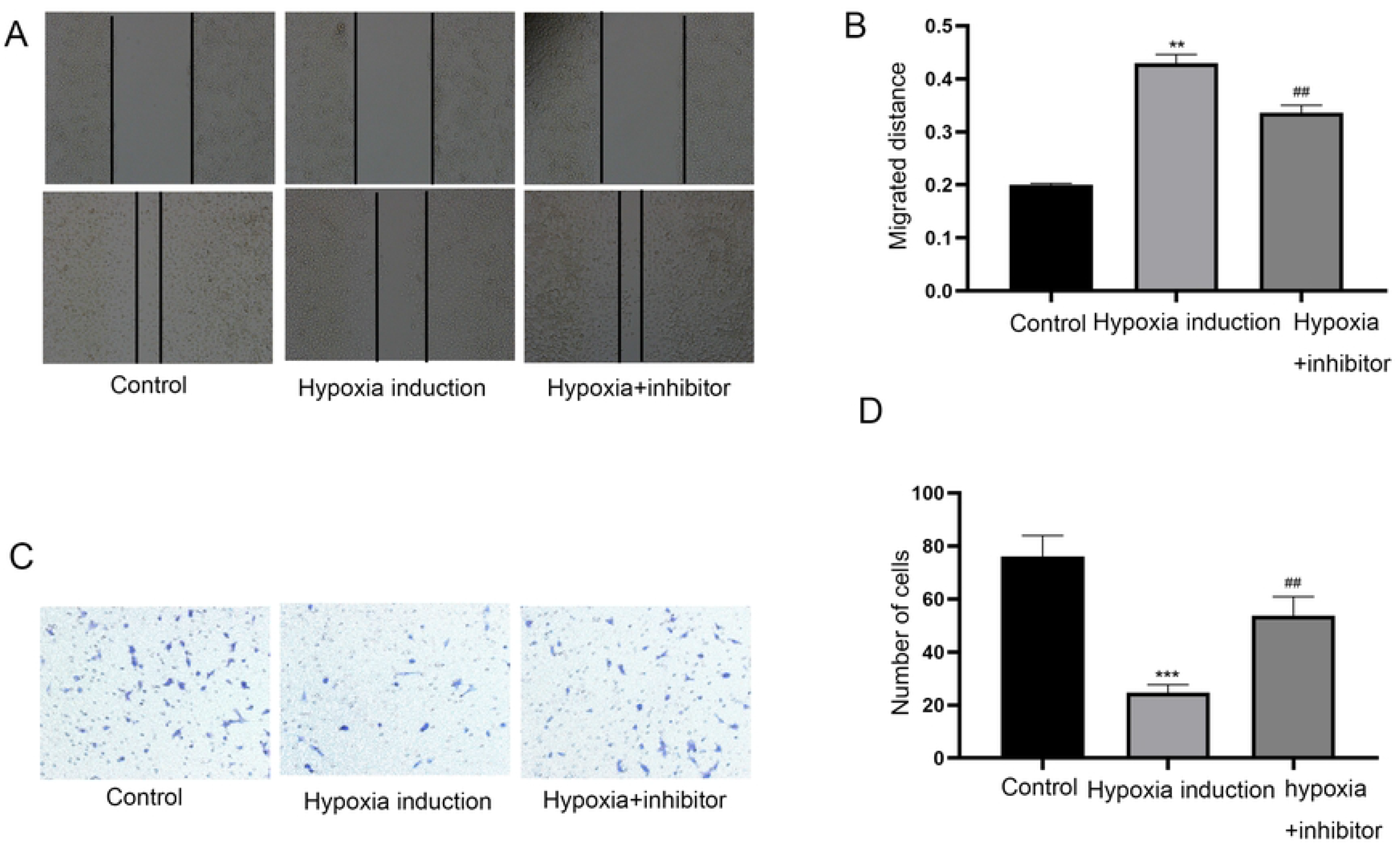
miR-195-3p inhibitor regains hypoxia-induced migration abilities of HUVECs. (**A, B**) Representative images of the scratch-wound healing assay under a microscope, the migrated distances of HUVECs were quantified. (**C, D**) The photos represent cells migrating on the bottom layer of the Transwell membrane. All photos were captured at 200× magnification. The total number of cells at each field was counted. The numbers of migrating cells are shown in the histograms (n=3) **p < 0.01, ***p < 0.001 vs. the control group. ##p < 0.01 vs. hypoxia induction for 6 h group. HUVECs, human umbilical vein endothelial cells.

### miR-195-3p inhibitor rescued the apoptosis of hypoxia injured HUVECs

To verify whether miR-195-3p inhibitor inhibits apoptosis, HUVECs transfected with miR-195-3p inhibitor for 48 h and subjected to hypoxia for 6 h were stained using TUNEL. Treatment of hypoxia injured HUVECs with miR-195-3p inhibitor significantly reduced the number of TUNEL-positive cells compared to the control cells (p < 0.0001) (Figure 5A, B). The expression levels of Bcl-2/Bax, cleaved caspase-3/caspase-3, and cytochrome C were also decreased in HUVECs transfected with miR-195-3p inhibitor (Figure 5C–F). These results suggest that the miR-195-3p inhibitor reduced the apoptosis of HUVECs induced by hypoxia.

**FIGURE 5.**
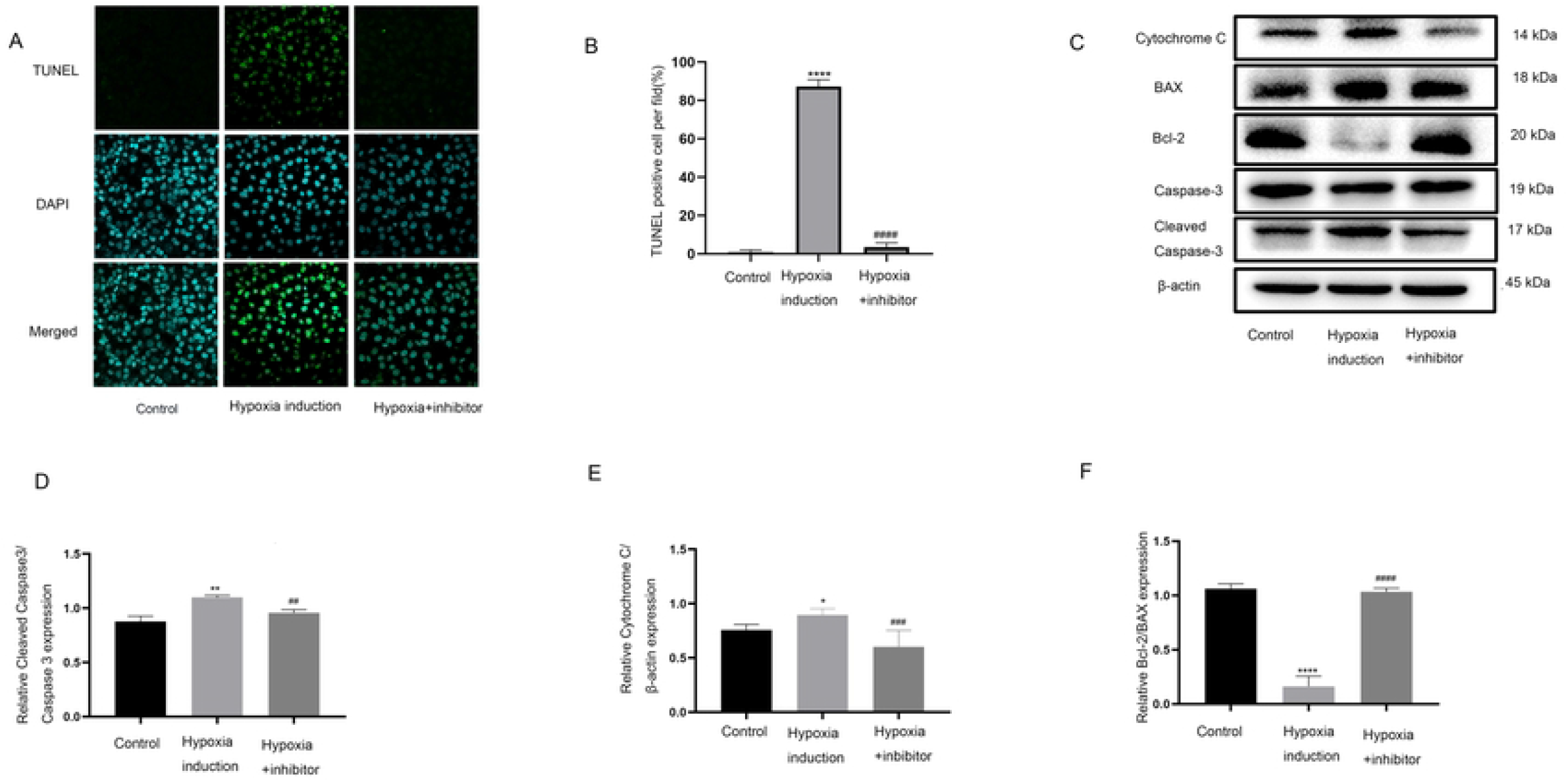
The miR-195-3p inhibitor attenuated hypoxia-induced HUVEC apoptosis. (**A**) Representative images of TUNEL staining of HUVEC showing the apoptotic cells (apoptotic cells stained in green and nucleus stained in blue with DAPI) Scale bar, 50 μm. (**B**) Statistical results of TUNEL-positive cells per field (n=3). (**C**) Western blot bands of control, hypoxia induction for 6 h, and hypoxia induction (6 h) + miR-195-3p. (**D–F**) Expression of cleaved caspase-3/caspase-3, Bcl-2/Bax, and cytochrome C were examined. β-actin served as an internal control. *p < 0.05, **p < 0.01, ***p < 0.001, ****p < 0.0001 vs. the control group. #p < 0.05, ##p < 0.01, ###p < 0.001, ####p < 0.0001 vs. hypoxia induction for 6 h group. HUVECs, human umbilical vein endothelial cells; TUNEL, terminal deoxynucleotidyl transferase dUTP nick end labeling; DAPI, 4’,6-diamidino-2-phenylindole.

We also found that treatment with miR-195-3p inhibitor could reverse the change in LC3B expression induced by hypoxia. The autophagic properties of HUVECs were restored (Figure 6).

**FIGURE 6.**
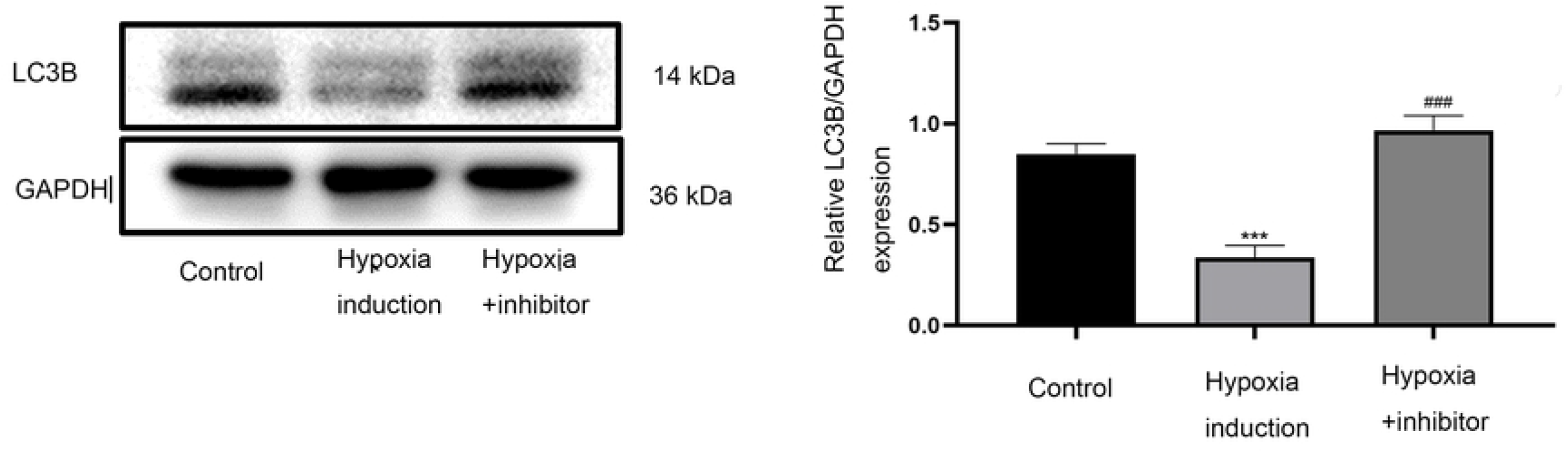
The miR-195-3p inhibitor regains hypoxia-induced HUVEC autophagy. Expression of LC3B in each group. miR-195-3p inhibitor can abrogate the changes in the expression of LC3B induced by hypoxia. Bands were quantified using Image J software. Each bar represents the mean ± standard deviation of three independent experiments. ****p < 0.0001 vs. the control group, ####p < 0.0001 vs. hypoxia induction for 6 h group.

## DISCUSSION

Herein, we found that inducing hypoxic injury in HUVECs resulted in a miR-195-3p-mediated decrease in cell proliferation, migration, and autophagy and an increase in cell apoptosis. A liposome-mediated transfection of HUVECs with miR-195-3p inhibitor downregulated miR-195-3p expression levels. Moreover, treatment with miR-195-3p inhibitor could partially reverse the hypoxic injury of HUVECs by enhancing cell proliferation, migration, and autophagy and reduced apoptosis.

In CHD, a blood clot or plaque causes reduced or insufficient blood flow to the heart muscles resulting in reduced oxygen and nutrition supply and subsequent myocardial damage(15). Angioplasty and surgical bypass are considered effective revascularization methods in most patients; however, stimulating the development of an effective collateral circulation has improved myocardial cell activity and prognosis in patients with severe CHD(16,17). The ECs are activated during collateral artery formation (arteriogenesis)(18), resulting in their proliferation, migration, and differentiation into tubular structures. The surrounding tissues wrap around the ECs to form vascular walls, and vascular networks coincide, initiating the collateral circulation(19–21). Researchers have a contrary opinion regarding the role of hypoxia in arteriogenesis. Even though hypoxia is thought to induce arteriogenesis, developing hypoxia is time-consuming. We, therefore, used HUVECs in our study to better understand the role of hypoxia on ECs. The “Anaerobic tank method” was used to establish the hypoxia model. The expression of HIF-1a in HUVECs(22,23) was analyzed to confirm the successful establishment of an in vitro hypoxia model. The experimental results confirmed that hypoxia injury weakened the proliferation, migration, and autophagy of HUVECs and promoted the apoptosis of HUVECs.

miRNAs regulate gene expression by binding to the 3’-UTR of the target mRNA, resulting in its translational repression(24,25). The role of miRNA in the development and progression of heart disease, especially CHD, has been described previously. An increased expression of miR-208b and miR-499 was reported during myocardial damage and is used to determine the severity of myocarditis(26). The genetic deficiency or pharmacological inhibition of miR-208a weakens stress-induced cardiac hypertrophy and remodeling(27). Therefore, identifying potential miRNAs involved in cardiovascular diseases and understanding the regulatory mechanism would be beneficial for the development of miRNA-based therapeutic interventions for heart disease. In recent years, the role of miR-195 in cardiovascular diseases has become increasingly prominent. Previous studies have shown that miR-195-5p inhibits the viability and proliferation and promotes apoptosis of HUVECs(28). In cardiomyocytes, miR-195 regulates the expression of mitochondrial apoptotic proteins and affects the survival of cardiomyocytes(29,30). However, the effect of miR-195-3p on hypoxia-induced HUVECs is rarely reported. This study showed that the expression of miR-195-3p increased significantly following induction of hypoxia in HUVECs. The hypoxic injury was associated with decreased proliferation and migration of the HUVECs and could be attributed to the direct or indirect effect of miRNA-195 on the G2/M phase of the target cell cycle (31). Bcl-2 has also been identified as a potential target for miRNA-195(32). A decrease in the Bcl-2/Bax ratio can activate caspase-3/9 resulting in apoptosis(33). Similar observations were made in this study, where the TUNEL-positive cells increased after hypoxic injury in HUVECs. A decrease in Bcl-2/Bax ratio and an increase in miR-195-3p, cleaved caspase-3/caspase-3 ratio, and cytochrome c levels confirmed hypoxia-induced apoptotic cell death in HUVECs.

Further treatment with a miR-195-3p inhibitor successfully downregulated its expression in HUVECs. Moreover, treatment with the inhibitor improved the migration and proliferation of HUVECs caused by hypoxia injury. A reversal in the characteristics of hypoxic injured HUVECs was observed upon treatment with the miR-195-3p inhibitor; number of TUNEL-positive cells were less, Bcl-2/Bax ratio increased, and apoptosis was significantly reduced, suggesting that miR-195-3p/Bcl-2 axis is involved in the mitochondrial apoptosis pathway after the HUVEC hypoxia injury. Autophagy refers to the process by which cells undergo self-degradation through a lysosome-dependent mechanism that has recently gained much attention in disease pathogenesis. LC3B is identified as a common autophagy marker(34,35). Previous studies have found a decreased expression level of LC3B protein following the hypoxic injury of HUVECs, resulting in inhibition of autophagy(36). Therefore, we studied the effect of the miR-195-3p inhibitor on autophagy in hypoxic HUVECs. Inhibition of miR-195-3p significantly increased the expression of LC3B, suggesting a role for miR-195-3p in regulating autophagy in HUVECs.

## CONCLUSION

In this study, we demonstrated the role of miR-195-3p in the hypoxic injury of HUVEC. Treatment with miR-195-3p inhibitor partially reversed the hypoxic injury of HUVEC, enhanced the migration, proliferation, and autophagy of cells, reduced apoptosis, and maintained the function of ECs under hypoxic conditions improving the activity of cardiomyocytes. Understanding the underlying molecular mechanism involved in miR-195-3p inhibitor-mediated rescue of hypoxic injury might provide new therapeutic strategies for treating CHD.

The limitations of our study are as followings: first of all, we only discussed Endothelial cell function, no further study of angiogenesis. Secondly,we only detected the change of miR-195-3p in vitro, but did not verify the changes in vivo. Due to the reason of time, we did not further carry out this study.

## Acknowledgements

Not applicable.

## Ethics statement

The experimental protocols were approved by the Ethics Committee of the Ethics of Experiments of Affiliated Hospital of Chengde Medical University, (Approval ID:CYFYLL 2022131).

## Availability of data and materials

The datasets used and/or analyzed during the present study are available from the corresponding author on reasonable request.

## Authors’ contributions

### Patient consent for publication

Not applicable

## REFERENCES

1. Sanada F, Taniyama Y, Muratsu J, et al. Gene-Therapeutic Strategies Targeting Angiogenesis in Peripheral Artery Disease. Medicines (Basel). Mar 30 2018;5(2)doi:10.3390/medicines5020031

2. Li L, Wang S, Wang M, Liu G, Yang Z, Wang L. miR-654-5p suppresses migration and proliferation of vascular smooth muscle cells by targeting ADAMTS-7. Cells Tissues Organs. 2022;doi:10.1159/000524677

3. Pulkkinen HH, Kiema M, Lappalainen JP, et al. BMP6/TAZ-Hippo signaling modulates angiogenesis and endothelial cell response to VEGF. Angiogenesis. Feb 2021;24(1):129–144. doi:10.1007/s10456-020-09748-4

4. Kanugula AK, Adapala RK, Jamaiyar A, et al. Endothelial TRPV4 channels prevent tumor growth and metastasis via modulation of tumor angiogenesis and vascular integrity. Angiogenesis. Aug 2021;24(3):647–656. doi:10.1007/s10456-021-09775-9

5. Zeng T, Zhang S, He Y, Liu Z, Cheng Q. MiR-361-5p promotes oxygen-glucose deprivation/re-oxygenation induced neuronal injury by negatively regulating SQSTM1 in vitro. Metab Brain Dis. Dec 2021;36(8):2359–2368. doi:10.1007/s11011-021-00845-x

6. van Rooij E, Sutherland LB, Liu N, et al. A signature pattern of stress-responsive microRNAs that can evoke cardiac hypertrophy and heart failure. Proc Natl Acad Sci U S A. Nov 28 2006;103(48):18255–60. doi:10.1073/pnas.0608791103

7. Tijsen AJ, van der Made I, van den Hoogenhof MM, et al. The microRNA-15 family inhibits the TGFβ-pathway in the heart. Cardiovasc Res. Oct 1 2014;104(1):61–71. doi:10.1093/cvr/cvu184

8. Zheng D, Ma J, Yu Y, et al. Silencing of miR-195 reduces diabetic cardiomyopathy in C57BL/6 mice. Diabetologia. Aug 2015;58(8):1949–58. doi:10.1007/s00125-015-3622-8

9. Hang P, Sun C, Guo J, Zhao J, Du Z. BDNF-mediates Down-regulation of MicroRNA-195 Inhibits Ischemic Cardiac Apoptosis in Rats. Int J Biol Sci. 2016;12(8):979–89. doi:10.7150/ijbs.15071

10. Zhang X, Ji R, Liao X, et al. MicroRNA-195 Regulates Metabolism in Failing Myocardium Via Alterations in Sirtuin 3 Expression and Mitochondrial Protein Acetylation. Circulation. May 8 2018;137(19):2052–2067. doi:10.1161/circulationaha.117.030486

11. Yang K, Zou Z, Wu Y, Hu G. MiR-195 suppression alleviates apoptosis and oxidative stress in CCl4-induced ALI in mice by targeting Pim-1. Exp Mol Pathol. Aug 2020;115:104438. doi:10.1016/j.yexmp.2020.104438

12. He X, Ji J, Wang T, Wang MB, Chen XL. Upregulation of Circulating miR-195-3p in Heart Failure. Cardiology. 2017;138(2):107–114. doi:10.1159/000476029

13. Aminzadeh A, Mehrzadi S. Melatonin attenuates homocysteine-induced injury in human umbilical vein endothelial cells. Fundam Clin Pharmacol. Jun 2018;32(3):261–269. doi:10.1111/fcp.12355

14. Yang NW, Kim JM, Choi GJ, Jang SJ. Development and evaluation of the quick anaero-system-a new disposable anaerobic culture system. Korean J Lab Med. Apr 2010;30(2):133–7. doi:10.3343/kjlm.2010.30.2.133

15. Lin M, Zhang Y, Niu Z, Chi Y, Huang Q. Transcriptomic responses of peripheral blood cells to coronary artery disease. Biosci Trends. Sep 19 2018;12(4):354–359. doi:10.5582/bst.2018.01078

16. Annibali G, Scrocca I, Aranzulla TC, Meliga E, Maiellaro F, Musumeci G. “No Reflow” Phenomenon: A Contemporary Review. J Clin Med. Apr 16 2022;11(8)doi:10.3390/jcm11082233

17. Lorentzen LG, Hansen GM, Iversen KK, Bundgaard H, Davies MJ. Proteomic Characterization of Atherosclerotic Lesions In Situ Using Percutaneous Coronary Intervention Angioplasty Balloons. Arterioscler Thromb Vasc Biol. Apr 21 2022:101161atvbaha122317491. doi:10.1161/atvbaha.122.317491

18. Chen M, Li X. Role of TRPV4 channel in vasodilation and neovascularization. Microcirculation. Aug 2021;28(6):e12703. doi:10.1111/micc.12703

19. Li B, Xian X, Lin X, et al. Hypoxia Alters the Proteome Profile and Enhances the Angiogenic Potential of Dental Pulp Stem Cell-Derived Exosomes. Biomolecules. Apr 14 2022;12(4)doi:10.3390/biom12040575

20. Jaskiewicz M, Moszynska A, Serocki M, et al. Hypoxia-inducible factor (HIF)-3a2 serves as an endothelial cell fate executor during chronic hypoxia. Excli j. 2022;21:454–469. doi:10.17179/excli2021-4622

21. Chen K, Wang Q, Liu X, Wang F, Yang Y, Tian X. Hypoxic pancreatic cancer derived exosomal miR-30b-5p promotes tumor angiogenesis by inhibiting GJA1 expression. Int J Biol Sci. 2022;18(3):1220–1237. doi:10.7150/ijbs.67675

22. Choudhry H, Harris AL. Advances in Hypoxia-Inducible Factor Biology. Cell Metab. Feb 6 2018;27(2):281–298. doi:10.1016/j.cmet.2017.10.005

23. Li M, Su Y, Gao X, Yu J, Wang Z, Wang X. Transition of autophagy and apoptosis in fibroblasts depends on dominant expression of HIF-1α or p53. J Zhejiang Univ Sci B. Mar 15 2022;23(3):204–217. doi:10.1631/jzus.B2100187

24. Paul S, Bravo Vazquez LA, Perez Uribe S, Roxana Reyes-Perez P, Sharma A. Current Status of microRNA-Based Therapeutic Approaches in Neurodegenerative Disorders. Cells. Jul 15 2020;9(7)doi:10.3390/cells9071698

25. Bartel DP. MicroRNAs: genomics, biogenesis, mechanism, and function. Cell. Jan 23 2004;116(2):281–97. doi:10.1016/s0092-8674(04)00045-5

26. Sluijter J, Deddens JC, Akker F, Hoogen P. Heart Failure in Chronic Myocarditis: A Role for microRNAs? Current Genomics. 2015;16(2):-.

27. Huang XH, Li JL, Li XY, et al. miR-208a in Cardiac Hypertrophy and Remodeling. Front Cardiovasc Med. 2021;8:773314. doi:10.3389/fcvm.2021.773314

28. Liao X, Zhou Z, Zhang X. Effects of miR1955p on cell proliferation and apoptosis in gestational diabetes mellitus via targeting EZH2. Mol Med Rep. Aug 2020;22(2):803–809. doi:10.3892/mmr.2020.11142

29. Gao CK, Liu H, Cui CJ, Liang ZG, Yao H, Tian Y. Roles of MicroRNA-195 in cardiomyocyte apoptosis induced by myocardial ischemia-reperfusion injury. J Genet. Mar 2016;95(1):99–108. doi:10.1007/s12041-016-0616-3

30. Xia H, Zhao H, Yang W, Luo X, Wei J, Xia H. MiR-195-5p represses inflammation, apoptosis, oxidative stress, and endoplasmic reticulum stress in sepsis-induced myocardial injury by targeting activating transcription factor 6. Cell biology international. 2021;doi:10.1002/cbin.11726

31. He JF, Luo YM, Wan XH, Jiang D. Biogenesis of MiRNA-195 and its role in biogenesis, the cell cycle, and apoptosis. J Biochem Mol Toxicol. Nov-Dec 2011;25(6):404–8. doi:10.1002/jbt.20396

32. Lin X, Zheng Z, Zhang Q, Zhang Z, An Y. Expression of miR-195 and its target gene Bcl-2 in human intervertebral disc degeneration and their effects on nucleus pulposus cell apoptosis. Journal of orthopaedic surgery and research. 2021;16(1):412. doi:10.1186/s13018-021-02538-8

33. Hu Y, Huang L, Shen M, et al. Pioglitazone Protects Compression-Mediated Apoptosis in Nucleus Pulposus Mesenchymal Stem Cells by Suppressing Oxidative Stress. Oxid Med Cell Longev. 2019;2019:4764071. doi:10.1155/2019/4764071

34. Fischer TD, Wang C, Padman BS, Lazarou M, Youle RJ. STING induces LC3B lipidation onto single-membrane vesicles via the V-ATPase and ATG16L1-WD40 domain. J Cell Biol. Dec 7 2020;219(12)doi:10.1083/jcb.202009128

35. Myszor IT, Sigurdsson S, Viktorsdottir AR, et al. The Novel Inducer of Innate Immunity HO53 Stimulates Autophagy in Human Airway Epithelial Cells. J Innate Immun. Jan 25 2022:1–16. doi:10.1159/000521602

36. Mai S, Muster B, Bereiter-Hahn J, Jendrach M. Autophagy proteins LC3B, ATG5 and ATG12 participate in quality control after mitochondrial damage and influence lifespan. Autophagy. Jan 2012;8(1):47–62. doi:10.4161/auto.8.1.18174

